# Hierarchical length and sequence preferences establish a single major piRNA 3’-end

**DOI:** 10.1101/2021.12.08.471772

**Authors:** Daniel Stoyko, Pavol Genzor, Astrid D. Haase

## Abstract

PIWI-interacting RNAs (piRNAs) guard germline genomes against the deleterious action of retroviruses and other mobile genetic elements. How piRNAs faithfully discriminate between self and non-self to restrict all mobile elements while sparing essential genes remains a key outstanding question in genome biology. PiRNAs use extensive base-pairing to recognize their targets and variable 3’ends could change the specificity and efficacy of piRNA silencing. Here, we identify conserved rules that ensure the generation of a single major piRNA 3’end in flies and mice. Our data suggest that the PIWI proteins initially define a short interval on pre-piRNAs that grants access to the ZUC-processor complex. Within this Goldilocks zone, the preference of the ZUC-processor to cut in front of a Uridine determines the ultimate processing site. We observe a mouse-specific roadblock that relocates the Goldilocks zone and generates an opportunity for consecutive trimming by PNLDC1. Our data reveal a conserved hierarchy between length and sequence preferences that controls the piRNA sequence space. The unanticipated precision of 3’end formation bolsters the emerging understanding that the functional piRNA sequence space is tightly controlled to ensure effective defense.

**Key findings:**

1. Every piRNA has a single major 3’end
2. Length preference trumps sequence preference during 3’end formation.
3. Relocation of the Goldilocks zone generates opportunity for trimming of mouse piRNAs.

## INTRODUCTION

Target-recognition by complementary base-pairing places the sequences of the guide-RNAs at the center of all RNA silencing mechanisms [1]. In eukaryotes, short guide RNAs associate with Argonaute proteins to form RNA-induced silencing complexes (RISC) that defend against viruses, regulate gene expression and protect genome integrity [2]. Within RISC, the sequence of the guide RNA determines target specificity while the Argonaute partner executes silencing mechanisms resulting in transcriptional and post-transcriptional restriction. The base-pairing interactions of guide RNAs and their targets follow common themes [3]. Universally, the most 5’ nucleotide of the guide RNAs is buried within the Argonaute protein and not available for base-pairing [4]. The following nucleotides 2-7 are the first to engage with a target [5, 6]. This ‘seed region’ dictates target specificity in microRNA (miRNA) silencing and provides a framework for studying other RNA silencing pathways [1]. Additional base-pairing across nucleotide 10 and 11 of the guide RNA is required for target-RNA cleavage by the intrinsic nuclease activity of Argonaute proteins [7, 8]. In contrast to miRNA-guided silencing, target recognition by PIWI-interacting RNAs (piRNAs) requires extensive base-pairing along the 3’ half of the guide RNA and potentially generates more specific and stable piRNA:target pairs [6].

PiRNAs and their PIWI protein partners silence endogenous retroviruses and other mobile genetic elements (transposons) to protect the integrity of germline genomes [9–11]. Mutants of key piRNA pathway genes universally result in the sterility of the animal. PiRNAs comprise millions of non-conserved sequences. How these diverse small RNAs faithfully restrict all transposons while avoiding potentially deleterious off-target effects remains unknown. Thousands of different piRNAs can be parsed from a single long precursor transcript. Processing is initiated by an endonucleolytic cut. Either the ZUC-processor complex or piRNA-guided slicing generates a 5’ monophosphorylated fragment that can be loaded onto a PIWI protein to form a pre-piRNA complex [11]. Consecutively, 3’end maturation is suggested to take place on the pre-assembled PIWI-pre-piRNA complex. The ZUC-processor complex generates a 3’ cut that releases the PIWI-piRNA complex from a potentially kilobases-long precursor [12–15]. Following this second endonucleolytic cut, a 3’-5’ exonuclease trims the 3’end to its mature length (**Fig. 1A**). This cut-n’-trim mechanism of piRNA 3’end formation has originally been suggested based on the observation that each PIWI protein associates with piRNAs of distinct lengths [16]. The footprint hypothesis suggests that 3’ends of piRNAs are determined by the physical impression of their PIWI-protein partner. Trimmers were identified in flies, silkworms, and mice [17–22]. PNLDC1, a 3’-5’ exonuclease related to Poly(A)-Specific Ribonuclease (PARN) trims pre-piRNAs to their mature length and is required for fertility in mice [17–19, 23]. An unrelated exonuclease, Nibbler (Nbr) trims some piRNAs, and select miRNAs in flies, but is dispensable for piRNA function [21, 22]. Fly piRNAs exhibit a preference for Uridine just after the mature 3’end (+1 position) corresponding to the signature of the ZUC-processor complex [12, 13, 24]. A +1U preference has also been observed for untrimmed pre-piRNAs in mouse [25], and additional sequence preferences were found for ZUC-mediated 3’end formation in silkworm cell extracts [26]. Taken together, these studies suggest that 3’end processing is guided by sequence preferences rather than length constraints. Here, we systematically probe how length and sequence preferences integrate into the 3’end decision and identify a conserved hierarchy of events. Our study elucidates an unanticipated precision in 3’end formation that generates a single major 3’end according to hierarchical length and sequence decisions.

**Figure 1.**
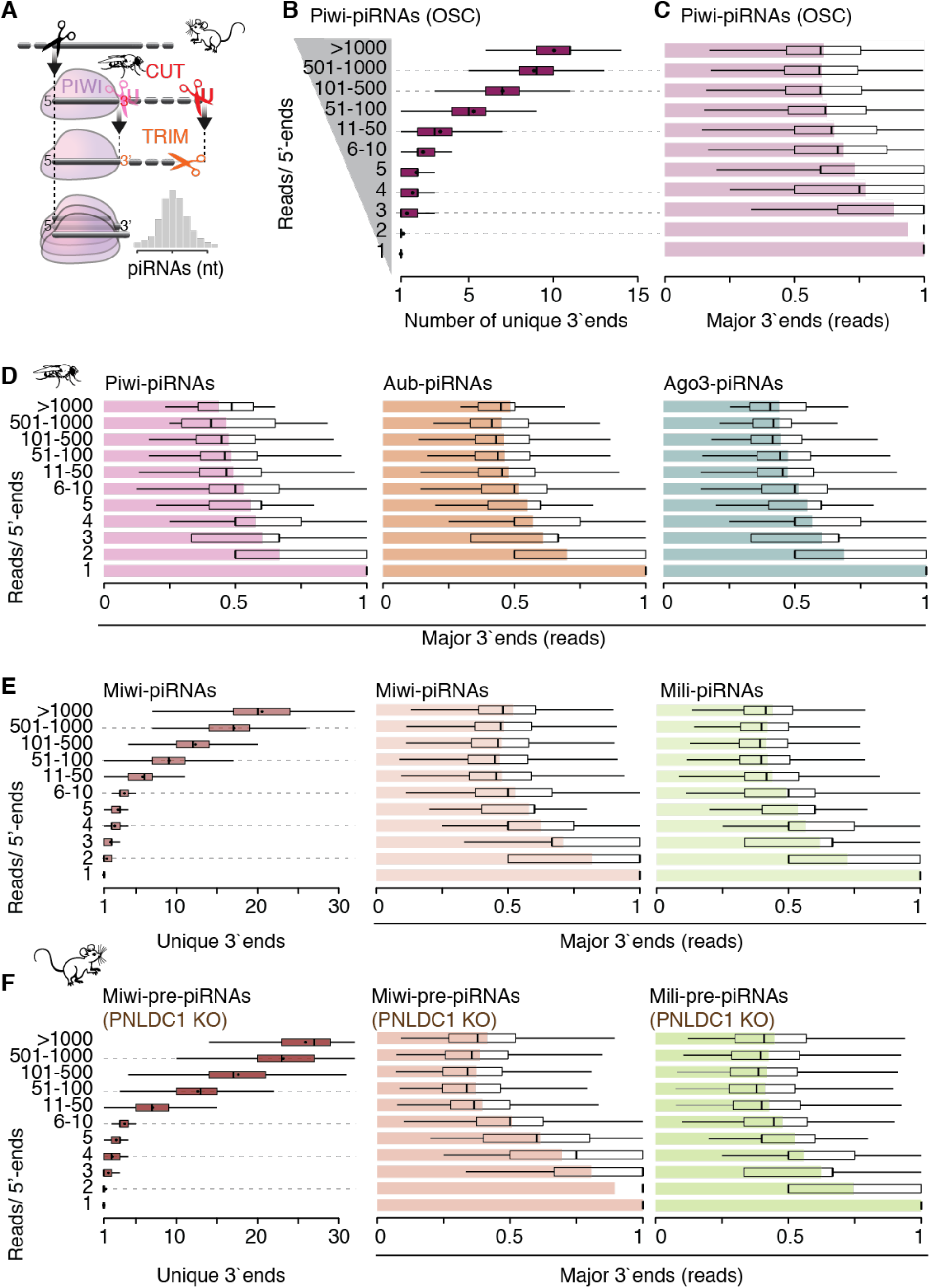
PiRNAs have a single major 3’-end. **(A)** The cut-n’-trim model for piRNA 3’end formation proposes an initial endonucleolytic cut that generates a substrate for 3’-to-5’ exonucleolytic trimming. The resulting length distribution of piRNAs are suggested to resemble a footprint of their bound PIWI protein partner. **(B)** Piwi-piRNAs that share the same 5’-end can have a number of different 3’-ends. Uniquely mapping Piwi-piRNAs from OSC (18-32 nt long) were grouped by the number of reads with a shared 5’-end. The number of different 3’-ends per 5’-end is depicted for each group. **(C and D)** Most of the reads associated with the same 5’-end have a single preferred 3’-end. For each 5’-end, the most abundant 3’-end was identified (referred to as “major”), and its contribution to the abundance of the 5’-end was calculated. The mean contribution is shown as bar plot while quartiles are depicted as a boxplot. Only uniquely mapping piRNAs that are 18-32nt long were used. **(E and F)** In mouse, both mature and untrimmed piRNAs can have many possible 3’-ends, however, only one contributes to approximately half of the abundance. Uniquely mapping reads were used for analyses (19-50nt). Data sets are publicly available (GSE156058; GSE83698; PRJNA421205).

## RESULTS

### Every piRNA has a single major 3’end

The length of piRNAs varies from ∼23-29nt in flies and ∼23-35nt in mice [9–11]. If piRNAs with the same 5’-end had a size distribution reflective of the total piRNA population, we would expect every 5’-end to be accompanied by multiple 3’-ends with the most prevalent represented by 20-30% of the small RNAs (**Fig. 1A**). To test this hypothesis, we first counted the total number of diverse 3’ends for each piRNA 5’end. To account for sequencing depth, we grouped piRNA 5’ends by read count reflecting the representation in small RNA molecules. For the most abundant Piwi-piRNAs in ovarian somatic sheath cells (OSC), with more than 1000 reads originating from the same 5’position, we observed on average ten different 3’ends (**Fig. 1B**). However, more than half of the reads were represented by a single major 3’end (**Fig. 1C**). We observed similar results for Piwi-piRNAs in fly ovaries, and Aubergine(Aub)- and Argonaute-3(Ago3) associated piRNAs, where more than half of all reads associated with a specific 5’end shared the same 3’end (**Fig 1D and Fig S1**) (ovary data [21]). Next, we analyzed available data for MIWI- and MILI-bound piRNAs in primary spermatocytes [25]. Like for fly piRNAs, we observed many possible 3’ ends for mature mouse piRNAs, but one of them prevailed (**Fig. 1E**). For highly represented MIWI-piRNAs (>1000 reads/ 5’end) we counted on average 21 different 3’ends. However, a single major end accounted for about half the reads, while the other half was represented by 20 different ends. Finally, we looked at murine pre-piRNAs that associated with MIWI or MILI in the absence of the trimming exonuclease (PNLDC1) (data [25]). Like mature piRNAs, pre-piRNAs had many possible 3’ends, but a single major end was represented by about half of all reads (**Fig. 1F**). Taken together, our data reveal that each piRNA has a single major 3’end.

### Major 3’ends emphasize length preferences specific for each PIWI protein

We next characterized the length preferences of major and minor 3’ends. The prevalence of a single major piRNA species associated with each 5’end eliminates 3’end variability as a reason for the broad length distribution of piRNA populations. It also suggests that individual major piRNAs (piRNAs that correspond to major 3’ends) are of different lengths, as no individual length accommodates more than half of all reads. To compare the length profiles of major (M) and minor (m) 3’ends, we normalized read counts within each group and plotted their distribution (**Fig. 2**).

**Figure 2.**
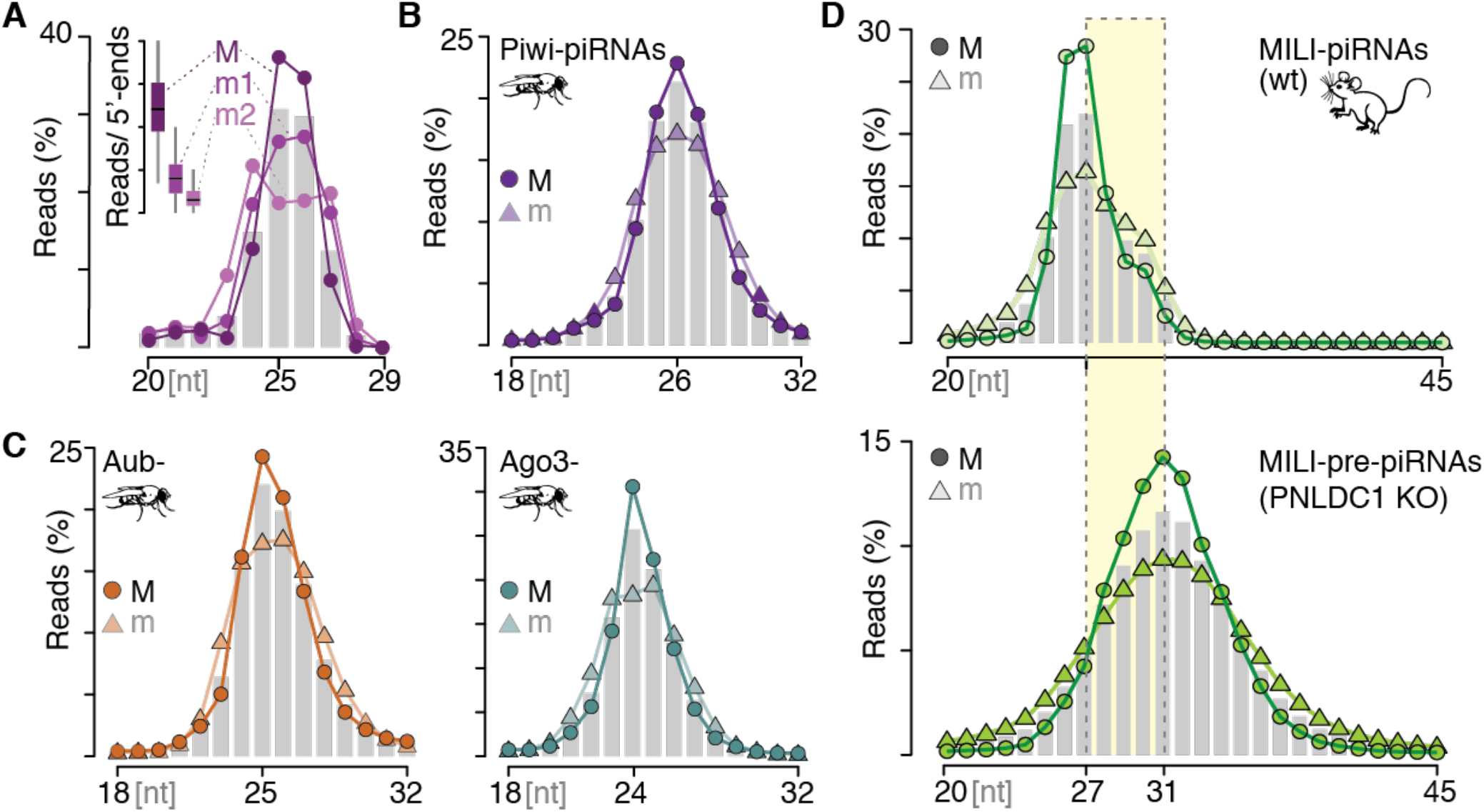
Major 3’ends reinforce specific length preferences. **(A)** Piwi-piRNAs from OSC that correspond to major 3’-ends have an increased preference to be 25 or 26 nucleotides (nt) long. PiRNAs with the most abundant (major; M), the second most abundant (‘first minor end’; m1), and the third most abundant (‘second minor end’; m2) 3’-ends were grouped, and their length distribution was calculated. Reads were normalized to the total reads witxhin each group. The bar plot depicts length distribution in nucleotides [nt]. The insert represents the contribution of Major (M) and minor (m1, m2) ends for each 5’-end. **(B and C)** Fly Piwi-, Aubergine (Aub-), and Argonaute3 (Ago3-) piRNAs with major 3’-ends have a preference to be 26, 25, and 24nt long respectively. The length distribution of major (M) and all minor (m) 3’-ends is normalized to the respective group. **(D)** Mature MILI-piRNAs that represent major 3’ends (M) have a stronger length preference than minor ends (m) in primary spermatocytes (SPI). MILI-pre-piRNAs are revealed in PNLDC1 knock-out (KO) spermatocytes (SPI) and exhibit a stronger length preference for major (M) compared to minor (m) sequences (data PRJNA421205).

Major Piwi-piRNAs are preferentially 25 or 26 nucleotides (nt) long in OSC (**Fig. 2A**). In contrast, the second most abundant end (m1; ‘first minor end’) shows less preference for a particular length, and the next end (m2, ‘second minor end’) starts to show a bimodal length distribution. In fly ovaries, major Piwi-piRNAs are preferentially 26nt long (**Fig. 2B**), while Aub- and Ago3-piRNAs peak at lengths of 25 and 24nt respectively (**Fig. 2C**). Minor piRNAs universally show less length preference. Interestingly, both, trimmed piRNAs and untrimmed pre-piRNAs show an increased length preference associated with their major ends in mouse spermatocytes (**Fig. 2D and S2A**). Major MILI-piRNAs peak at a length of 27nt while MIWI-piRNAs peak at 30nt. Their major untrimmed counterparts are preferentially four nucleotides longer, resulting in 31nt long pre-piRNAs associated with MILI and 34nt long MIWI-pre-piRNAs. To test whether this four-nucleotide difference represents the preference of individual trimming events, we identified piRNA:pre-piRNA pairs and calculated the 3’distance of their major ends. Our data show that major pre-piRNAs associated with MIWI and MILI are preferentially four nucleotides longer than their mature counterparts (**Fig. S2B**). The distinct difference of mature and pre-piRNAs in mouse suggests a consistent roadblock that prevents access for the ZUC-processor on the pre-piRNA. Overall, the reinforced length preferences of major piRNAs bolster the hypothesis that 3’ends are defined through the footprint of their associated PIWI protein.

### Major 3’cuts bolster a preference for Uridine following the cleavage site (+1U)

Next, we characterized sequence preferences at the major cut sites. The ZUC-processor complex establishes a preference for Uridine in the +1 position, and a +1U-bias has been observed for mature piRNAs in flies and untrimmed pre-piRNAs in mice [12, 13]. We observed a gradual decrease in +1U preference from the major (M) to the minor 3’ends (m1, 2) in Piwi-piRNAs (**Fig. 3A**). Universally, we observed a stronger +1U preference for Major (M) compared to minor (m) piRNAs associated with all three PIWI proteins in flies (**Fig. 3B**). Consistent with the idea that mature 3’ends in mouse are not guided by sequence preferences [13, 25], we did not observe a difference in +1U abundance between major and minor MILI- and MIWI-piRNAs in mouse spermatocytes (**Fig. 3C**). However, major pre-piRNAs showed a strong enrichment of +1U compared to minor 3’ends in PNLDC1 knockout mice (**Fig. 3D**). Overall, we observed an increased preference for Uridine one nucleotide downstream of major 3’ends when compared to minor ends, supporting the hypothesis that 3’ends are defined by sequence preference.

**Figure 3.**
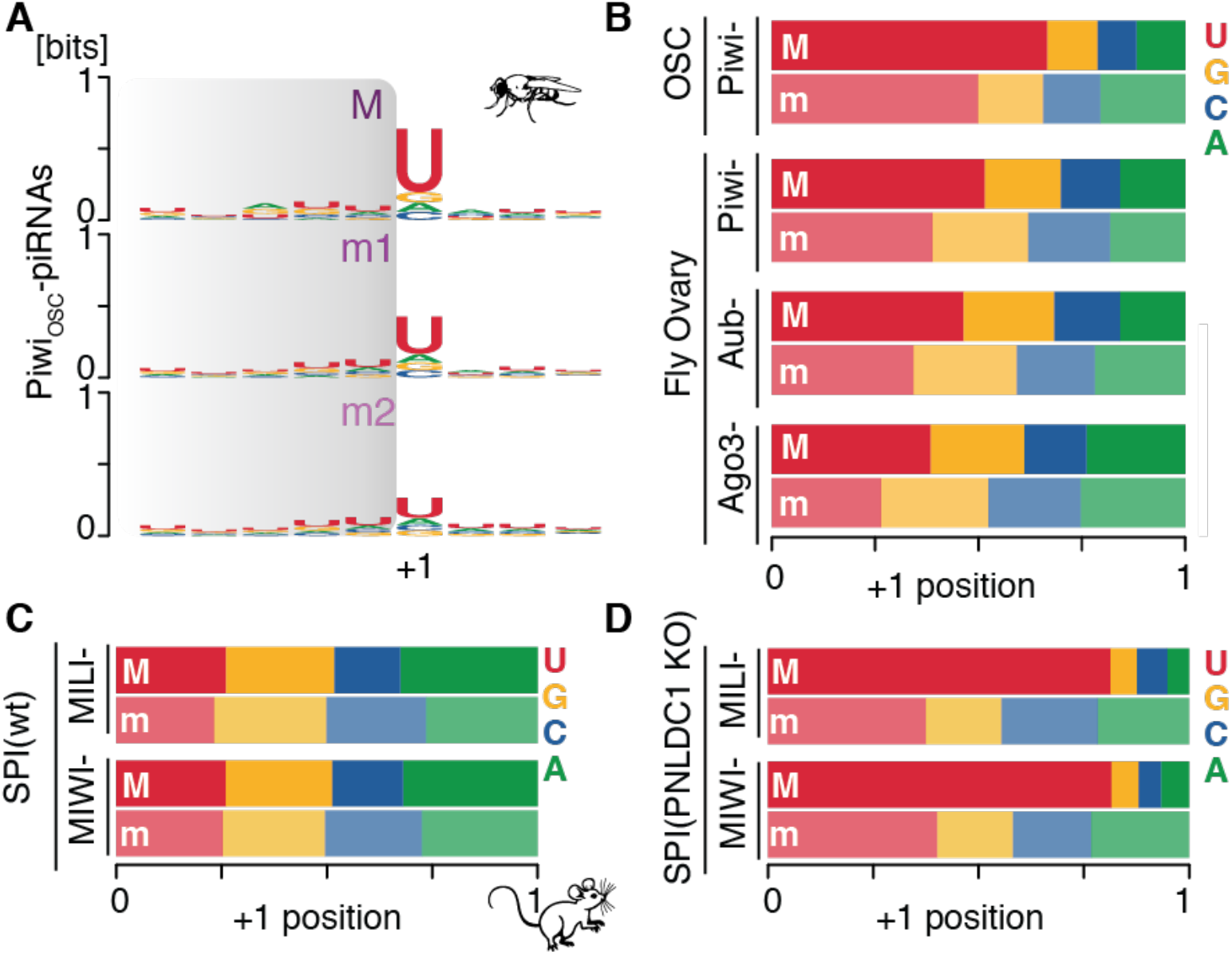
Major 3’-cuts show an increased preference for Uridine in the +1 position. **(A)** Piwi-piRNAs with major 3’-ends (M) have a stronger preference for Uridine in the +1 position (+1U) than minor 3’ends (m). Sequence preferences of the most abundant (major; M), the second most abundant (minor; m1), and the third most abundant (minor; m2) 3’-ends are depicted as sequence logo in bits. The observed part of the piRNAs is indicated in grey. The +1 position marks the nucleotide immediately following the piRNA 3’end. **(B)** Major 3’-ends increase the +1U preference for piRNAs associated with all three PIWI proteins in *Drosophila* ovaries. The nucleotide composition in the +1 position was calculated for major (M) and all minor (m) piRNA 3’-ends. **(C)** Neither major (M) nor minor (m) 3’ends of mature (trimmed) murine piRNAs exhibit a +1U preference. **(D)** The fraction of +1U following a major 3’end is doubled compared to minor 3’ends of untrimmed pre-piRNAs in mice. (Uniquely mapping reads were used for all panels).

### Length preference trumps sequence preference during 3’end formation

Our characterization of major piRNA 3’ends produced evidence for length and sequence preferences in flies and mice. Next, we aimed to identify how these preferences interact to define a single major 3’end. To this end, we probed the impact of Uridines at different positions relative to a preferred length (**Fig. 4**). Piwi-piRNAs are preferentially 25 and 26 nt long. The presence of a Uridine one nucleotide downstream (position 26 and 27) results in a repositioning of 3’ends, and close to half of all piRNAs now end 5’ of this Uridine (**Fig. 4A**). Uridines at positions outside of this preferred distance from the 5’end show no major effect on 3’end positioning. In the absence of any Uridine in a five-nucleotide window surrounding position 26, 3’ ends remain at positions 25 and 26 irrespectively. The effect of Uridine on the position of piRNA 3’ends can also be observed by calculating a Z-score for 3’ends across a local genomic interval, and when we restrict the nucleotide composition surrounding the Uridine to Adenine (A), Cytosine (C) or Guanosine (G) (**Fig. 4B**). Finally, we ask what happens in the presence of Poly-Uridines. For Piwi-piRNAs we observe a preference to position the 3’ end just before the last of two or three Uridines (**Fig. 4C**). This preference might contribute to the observed 3’signature of ZUC-dependent piRNA processing in silkworm [26], and to the enrichment of UVV triplets following the 3’end of murine piRNAs [27]. We observe similar positional effects for Uridines for Piwi-, Aub- and Ago3-piRNAs in fly ovaries (**Fig. S3**) and pre-piRNAs in mice (**Fig. 4D**). Our data suggest that PIWI proteins establish a Goldilocks zone for Uridines to impact 3’end processing.

**Figure 4.**
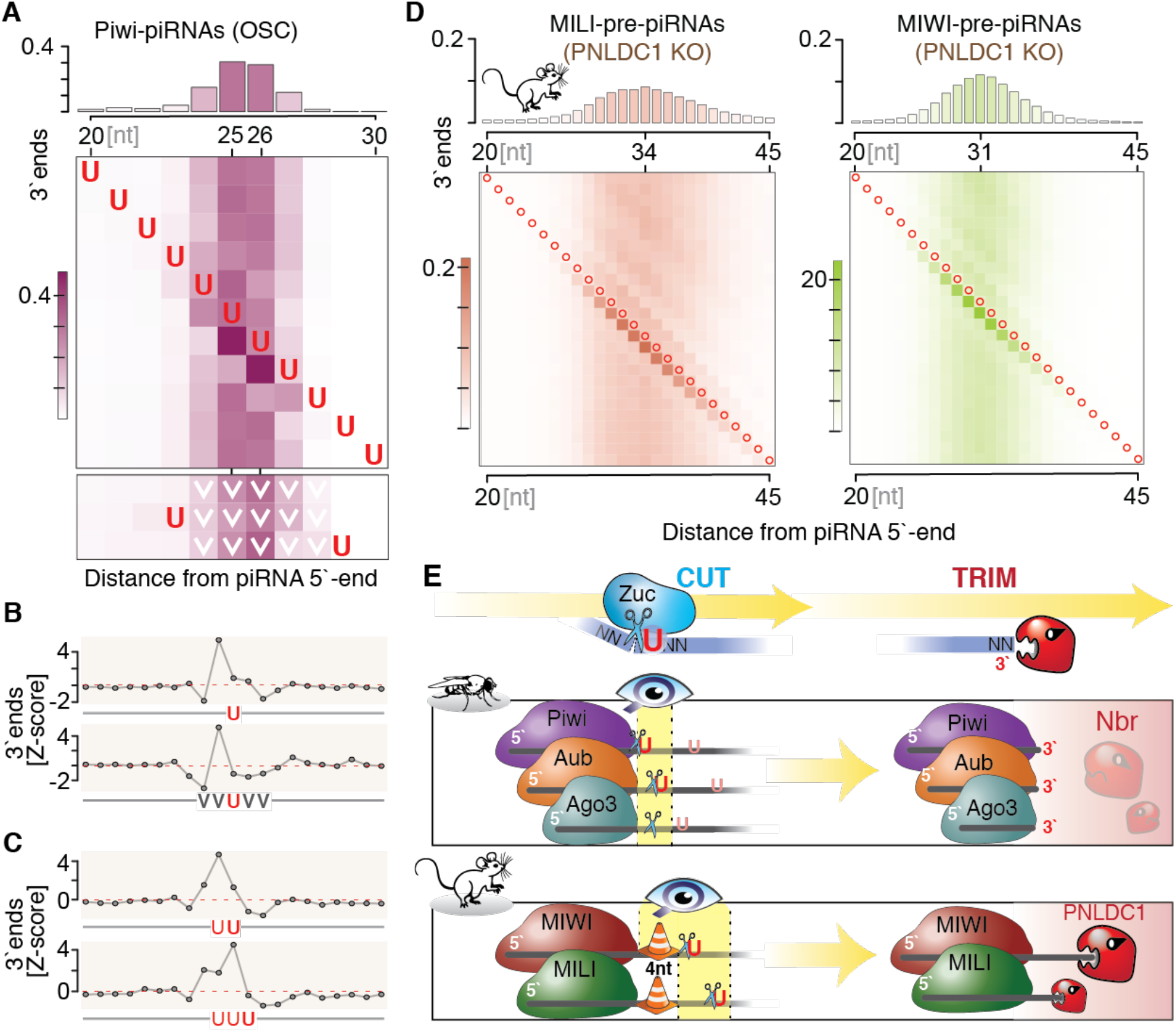
Length preference is predominant over +1U preference. **(A)** For Piwi-piRNAs (OSC), the 3’-end is defined by the location of Uridine (U) within a narrow window located at a fixed distance from the 5’-end. The top panel depicts the length distribution of Piwi-piRNAs for orientation. Each row of the heatmap indicates fractions of piRNA 3’ends when the position of a single Uridine (U) is fixed across the interval. The heatmap at the bottom shows fractions of 3’ends in the absence of Uridines (U) across a continuous five nucleotide region (row 1), and upon requirement for a Uridine (U) immediately 5’ or 3’ of the fixed non-U (V) stretch. (V is C, G or A). **(B)** PiRNA 3’-ends are more likely to occur before Uridines. Z-score was calculated over 51-nt window (central 21nt are depicted), with the Uridine (U) located in the center. **(C)** In the presence of a poly-Uridine sequence (UU or UUU), piRNA 3’-ends are preferentially located before the last Uridine. (Z-score calculated across a 51-nt window; the central 21nt are depicted.) **(D)** The presence of Uridines within an optimal length interval - the Goldilocks zone-shifts the preferred 3’ end of murine pre-piRNAs. Red circles indicate Uridines (U). (**E**) The revised model for piRNA 3’end formation shows a hierarchical integration of length and sequence preferences to generate a single major piRNA species. First PIWI determines a specific interval for access to the ZUC-processor complex on the pre-piRNA. Within this Goldilocks zone, the ZUC-processor preferentially cuts 5’ of a Uridine (U). A mouse-specific roadblock changes the position of the Goldilocks zone and leaves substrate for consecutive trimming by PNLDC1. The final major piRNA likely reflects a footprint of its associated PIWI protein.

Overall, a model emerges that integrates length and sequence preferences to establish a single major 3’end for piRNAs in flies and mice (**Fig. 4E**). Universally, length preferences trump sequence preferences in a hierarchical decision. In flies, the ZUC-processor complex seems to have access to the pre-piRNA immediately downstream of PIWI’s footprint, which does not leave much room for trimming after the initial cut. In contrast, ZUC’s visibility is shifted by a consistent distance on murine pre-piRNAs. After the initial cut, the roadblock seems to be lifted and PNLDC1 faithfully trims four nucleotides to establish the mature 3’end with major sequences that are determined by length preferences.

## DISCUSSION

Each PIWI protein establishes a distinct Goldilocks zone for 3’end processing on pre-piRNAs. Whether these specific footprints reflect the geometry of different PIWI proteins, post-translational modifications or function requires further studies. Length preferences could reflect the most stable conformation of PIWIs in complex with their guide RNAs or optimal (in)stability for effective silencing. Transcriptional silencing might benefit from increased stability of piRISC:target interactions and a longer resident time on chromatin. However, there is no simple relationship between length and silencing mode. In *Drosophila* the nuclear PIWI protein, Piwi, associates with the longest piRNAs, while in mouse and human the longest piRNAs are found in the cytoplasmic PIWIL1 (MIWI or HIWI). Cytoplasmic slicing of target transcripts by piRISC bares similarity to target cleavage by Argonaute complexes during RNA interference (RNAi), and additional base-pairing interactions are likely to impact turn-over kinetics [28]. Sequence, length, and modifications impact piRNA stability [29], and it would be interesting to know, if individual PIWI proteins have an intrinsic length optimum for their function. Finally, length determines the sequence space available for target interaction. In contrast to the well-characterized seed-recognition of miRNAs, seed-integrations of piRNAs with a target are weaker and compensated by extensive base-pairing across the piRNA length [1, 6]. Longer piRNAs have the potential for stronger interactions, and precision of a major 3’end might be required to control specificity. The notion that every piRNA has a single major 3’end contributes to our emerging understanding that the functional piRNA sequence space is regulated to ensure effect genome defense [30].

## MATERIALS and METHODS

### Data access and initial processing of sequencing data

A total of 12 datasets were analyzed in this study. OSC data were obtained from [GEO accession number: GSE156058]. FH-Piwi-IP dataset was generated by combining three replicates with the following SRA accession numbers: SRR12430857, SRR12430858, SRR12430859. Drosophila ovary data were obtained from [GSE83698]. SRA accession numbers are as follows: fly-Piwi-IP = SRR3715419, fly-Aub-IP = SRR3715420, fly-Ago3-IP = SRR3715421. Mouse data were obtained from [NCBI BioProject accession number PRJNA421205]. Datasets were initially processed as previously described [30]. All computational code is available in ‘Supplemental Code’.

### Counting numbers of unique 3’-ends: (Fig 1B, 1E, 1F, S1A

Uniquely mapping piRNAs (NH = 1) were collapsed by sequence while retaining their abundance information (# of reads per unique sequence). The following sequences were collapsed again by the coordinate of their 5’-end while retaining the total read abundance and the number of unique sequences that share the 5’-end coordinates. These 5’-ends were subsequently grouped into discrete bins based on their total read abundance (bins shown on y-axis). Amounts of unique sequences sharing 5’-ends were depicted for each bin using geom_boxplot() function from the ggplot package for R.

### Defining the “major 3’-end” and determining its contribution: (Fig 1C, 1D, 1E)

Uniquely mapping piRNA sequences (NH = 1) were grouped by their 5’-end coordinates. For each 5’-end, the most abundant sequence was identified as “major end”. Ties for the title of “major end” were solved in a random fashion to avoid biases. The 5’-ends were subsequently grouped into discrete bins based on their total read abundance (bins shown on y-axis). For each bin, the contribution of the “major end” was determined by dividing the abundance of the “major end” by the total abundance of all sequences sharing the respective 5’-end. The mean value for each bin was depicted as bar plot using geom_barplot() while the distribution of contribution values within each bin was depicted as a boxplot using geom_boxplot() from the ggplot package.

### Ranking 3’-ends: (Fig 2A)

Uniquely mapping piRNA sequences (NH = 1) were once again grouped by their 5’-end coordinates. For each 5’-end, the most abundant sequence (with major 3’-end) was identified as “M”, the second most abundant sequence was identified as “m1” and the third as “m2”. The contribution of these sequences to the abundance of the 5’-end was calculated in the same fashion as previously. The distribution of these contribution values was plotted as a boxplot in the insert of Fig 2A. The size distributions of “M”, “m1”, and “m2” were calculated by counting the number of reads at different sizes and normalizing by the total number of reads in “M”, “m1”, or “m2” bins.

### Size distribution of the “major 3’-end”: (Fig 2B–D, S2A)

For each 5’-end, a “major end” was identified in the same manner as described previously. Size distribution of all “major ends” was determined by counting the number of “major end” reads at different lengths and normalizing by the total number of “major end” reads. Same analysis was performed on “minor ends” whereby the “minor ends” are any sequences that were not identified as “major ends”. The size distribution of the entire library was depicted as bar plot using the ggplot package for R.

### Distance between major 3’-ends in trimmed vs mature piRNAs: (Fig S2B)

The major 3’-end was defined for uniquely mapping piRNA sequences (NH = 1) in WT and PNLDC KO samples. Only major 3’-end containing sequences whose 5’-end was present in both datasets were kept. The resulting sequences in the WT sample were subsequently paired with the corresponding sequence (of the same 5’-end) in PNLDC-KO sample. Only pairs where the PNLDC-KO sequence was longer than the WT sequence were kept. The nucleotide length difference between these two sequences was determined and depicted as a bar plot using the ggplot package.

### Determining the composition of the +1 nucleotide: (Fig 3)

Uniquely mapping piRNA sequences (NH = 1) were categorized into “major ends” and “minor ends” in a manner described previously. The identity of the nucleotide immediately after the 3’-end of piRNAs (referred to as the +1 nucleotide) was obtained from the corresponding genome (dm6 for fly and mm10 for mouse). The composition of this nucleotide was depicted as a stacked bar plot using the ggplot package. Sequences logos were generated using the ggseqlogo package for R.

### Sequence context specific 3’-end distribution: (Fig 4A, 4D)

Uniquely mapping piRNA sequences (NH = 1) were grouped into non-mutually exclusive bins based on the sequence context downstream of the piRNA 5’-end. The distribution of 3’-ends for each sequence context was determined by counting the number of reads at different sizes and normalizing by the total amount of reads within the sequence context. This distribution was plotted as a heatmap using the geom_tile() function within the ggplot package.

### Z-score for 3’-ends: (Fig 4B, 4C)

Genomic sequence immediately downstream and upstream of all piRNA (uniquely mapping only) 3’-ends was isolated. A sequence context of interest (ex: “UUU”) was searched for within a 51nt window surrounding the 3’-end. If found, information regarding the position of this sequence context in respect to the 3’-end was recorded. Data from all successfully located sequence contexts were collated to determine total numbers of piRNA 3’-ends at given distances from the sequence context of interest. The total number of piRNA 3’-ends for each position was converted to a Z-score by determining the number of standard deviations from the mean (within the full 51nt window).

### DATA ACESS

All sequencing data are publicly available: GSE156058[30], GSE83698[21], PRJNA421205[25].

## ACKNOWLEDGEMENTS

We thank M Hafner for critical comments on the manuscript, all members of the Haase and Hafner labs for helpful discussions and E He (NIH Medical Arts) for help with the model. This work would not have been possible without the support of the NIH high-performance computing group and funding by the intramural research program of the NIDDK.

## AUTHOR CONTRIBUTIONS

D.S. designed and performed all experiments with help from P.G.. A.D.H. and D.S. wrote the manuscript.

## Supplemental igures and figure legends (S1-3)

**Supplemental Figure 1.**
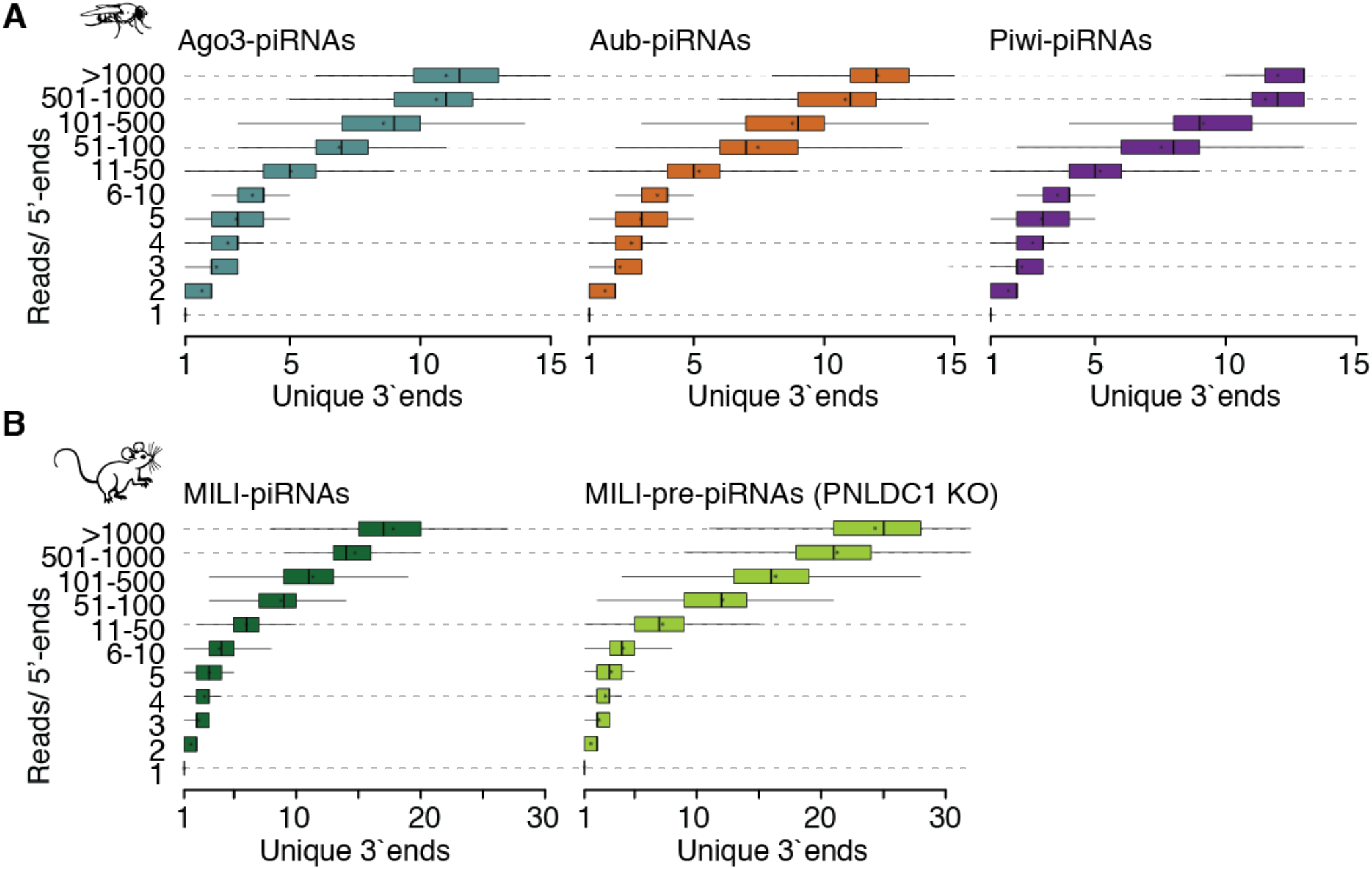
Fly and mouse piRNAs have a many possible 3’-ends. **(A)** *Drosophila* piRNAs with the same 5’-end have a number of different 3’-ends. Uniquely mapping piRNAs (18-32nt in length) were grouped by the number of reads sharing their respective 5’-ends. Number of unique 3’-ends observed per 5’-end is depicted for each group. **(B)** Both mature and untrimmed MILI-piRNAs can have several possible unique 3’-ends. (Uniquely mapping piRNAs were used.)

**Supplemental Figure 2.**
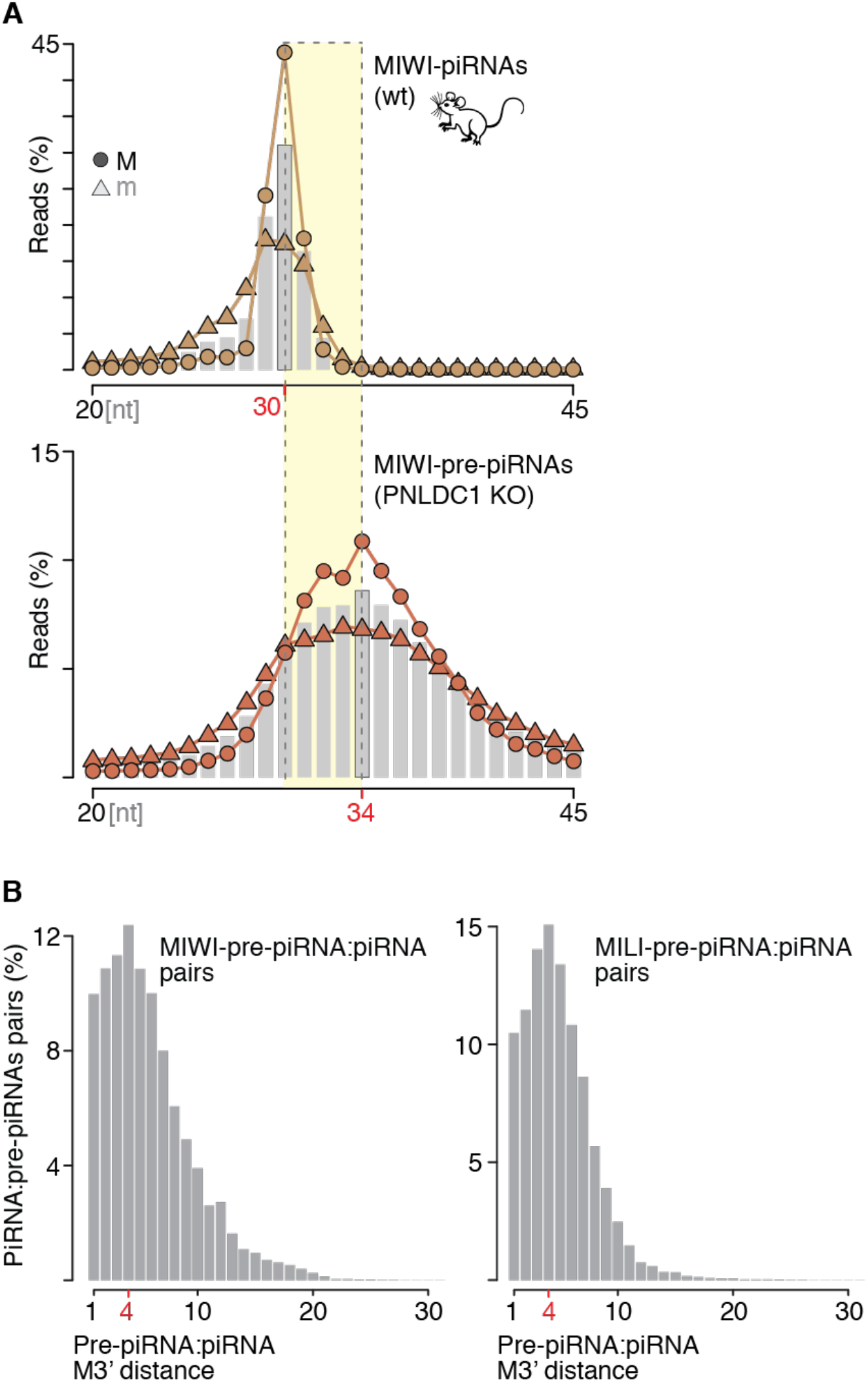
Major 3’-ends of murine piRNAs have specific length preferences that are four nucleotides (4nt) apart. **(A)** Both, trimmed and untrimmed MILI-piRNAs with major 3’-ends have distinct length preferences at 30nt and 34nt respectively. The length distribution of major (M) and all minor (m) 3’-ends is normalized to the respective group. Bar plot depicts length distribution of the entire library. **(B)** Untrimmed murine piRNAs are preferentially 4nt longer than their mature counterparts. Untrimmed-piRNAs were paired with trimmed-piRNAs that have the same 5’-end. The distance between the major 3’-ends of the pair was calculated and scaled by the number of piRNAs in the pair.

**Supplemental Figure 3.**
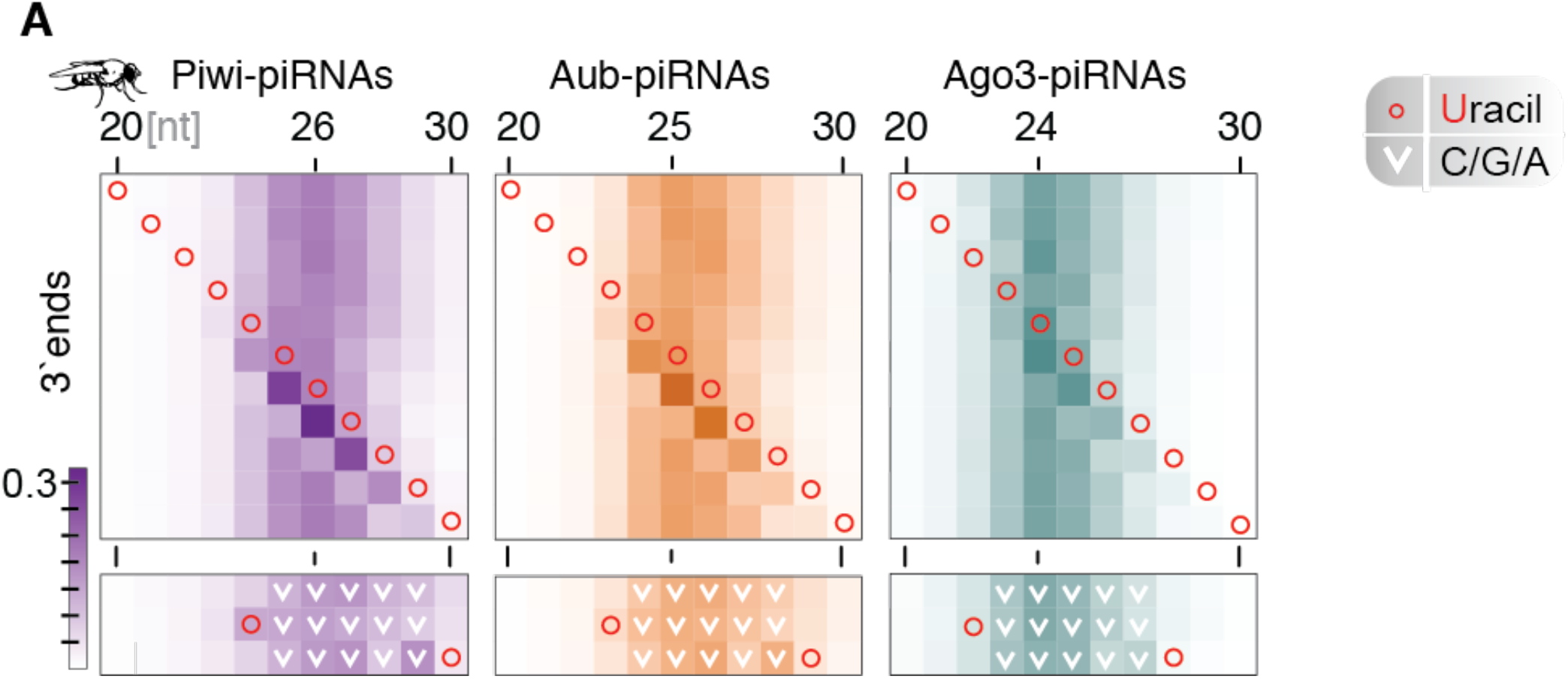
The presence of Uridines (U) affects the position of 3’ends only within an optimal distance from the piRNA 5’end. **(A)** In fly ovary piRNAs, the 3’-end is defined by the location of Uridine within a narrow window located at a fixed distance from the 5’-end. Rows depict distribution of 3’-ends for piRNAs where a Uridine (red circle) or absence of Uridine (V) is present at a certain distance from the 5’-end.

